# Bicarbonate Activation of Monomeric Photosystem II-PsbS/Psb27 Complex

**DOI:** 10.1101/2022.01.13.476245

**Authors:** Andrea Fantuzzi, Patrycja Haniewicz, Domenica Farci, M. Cecilia Loi, Keunha Park, Claudia Büchel, Matthias Bochtler, A. William Rutherford, Dario Piano

## Abstract

In thylakoid membranes, Photosystem II monomers from the stromal lamellae contain the subunits PsbS and Psb27 (PSIIm-S/27), while Photosystem II monomers from granal regions (PSIIm) lack these subunits. Here, we have isolated and characterised these two types of Photosystem II complexes. The PSIIm-S/27 showed enhanced fluorescence, the near-absence of oxygen evolution, as well as limited and slow electron transfer from Q_A_ to Q_B_ compared to the near-normal activities in the granal PSIIm. However, when bicarbonate was added to the PSIIm-S/27, water splitting and Q_A_ to Q_B_ electron transfer rates were comparable to those in granal PSIIm. The findings suggest that the binding of PsbS and/or Psb27 inhibits forward electron transfer and lowers the binding affinity for the bicarbonate. This can be rationalized in terms of the recently discovered photoprotection role played by bicarbonate binding via the redox tuning of the Q_A_/Q_A_^•−^ couple, which controls the charge recombination route, and this limits chlorophyll triplet mediated ^1^O_2_ formation (Brinkert K et al. (2016) Proc Natl Acad Sci U S A. 113(43):12144-12149). These findings suggest that PSIIm-S/27 is an intermediate in the assembly of PSII in which PsbS and/or Psb27 restrict PSII activity while in transit, by using a bicarbonate-mediated switch and protective mechanism.

**One sentence summary:** A photosystem II monomer with PsbS and Psb27 as additional subunits, is inactive as isolated but activated by bicarbonate, and is attributed to be a late-stage intermediate in photoassembly.

## Introduction

Oxygenic photosynthesis is a light-driven biochemical process providing the biosphere with organic carbon, energy, and molecular oxygen (Johnson, 2016). In higher plants, this process takes place in the chloroplast, a specialized organelle consisting of outer and inner membranes forming a network of photosynthetic membranes named thylakoids (Albertsson, 2001; Vothknecht and Westhoff, 2002). The protein composition of the different portions of these membranes are distinct, showing segregation of the photosystems. Photosystem II (PSII) is mainly present in the appressed granal regions, while Photosystem I (PSI) is found in the non-appressed regions of the granal margins and in the stromal lamellae (Andersson and Anderson, 1980). Dynamic responses to various environmental factors have been shown to change the ultrastructure and composition of the membranes and photosystems (Ruban and Johnson, 2015; Kirchoff, 2019).

In higher plants, the organization of thylakoid membranes also reflects the partition of different kinds of PSII, which are assembled and repaired in the stromal lamellae, while the fully functional PSII complexes are located in the grana (Andersson and Anderson, 1980; Danielsson et al., 2006) (Fig. 1). The small fraction of PSII complexes that are found in stromal lamellae are mainly PSII monomers (PSIIm) and a series of smaller assembly intermediates. In contrast, the grana are dominated by PSII dimers that can form a range of complexes with chlorophyll antenna proteins, including Light Harvesting Complex II (LHCII), forming PSII-LHCII (Danielsson et al., 2006; Watanabe et al., 2009; Haniewicz et al., 2013).

**Figure 1:**
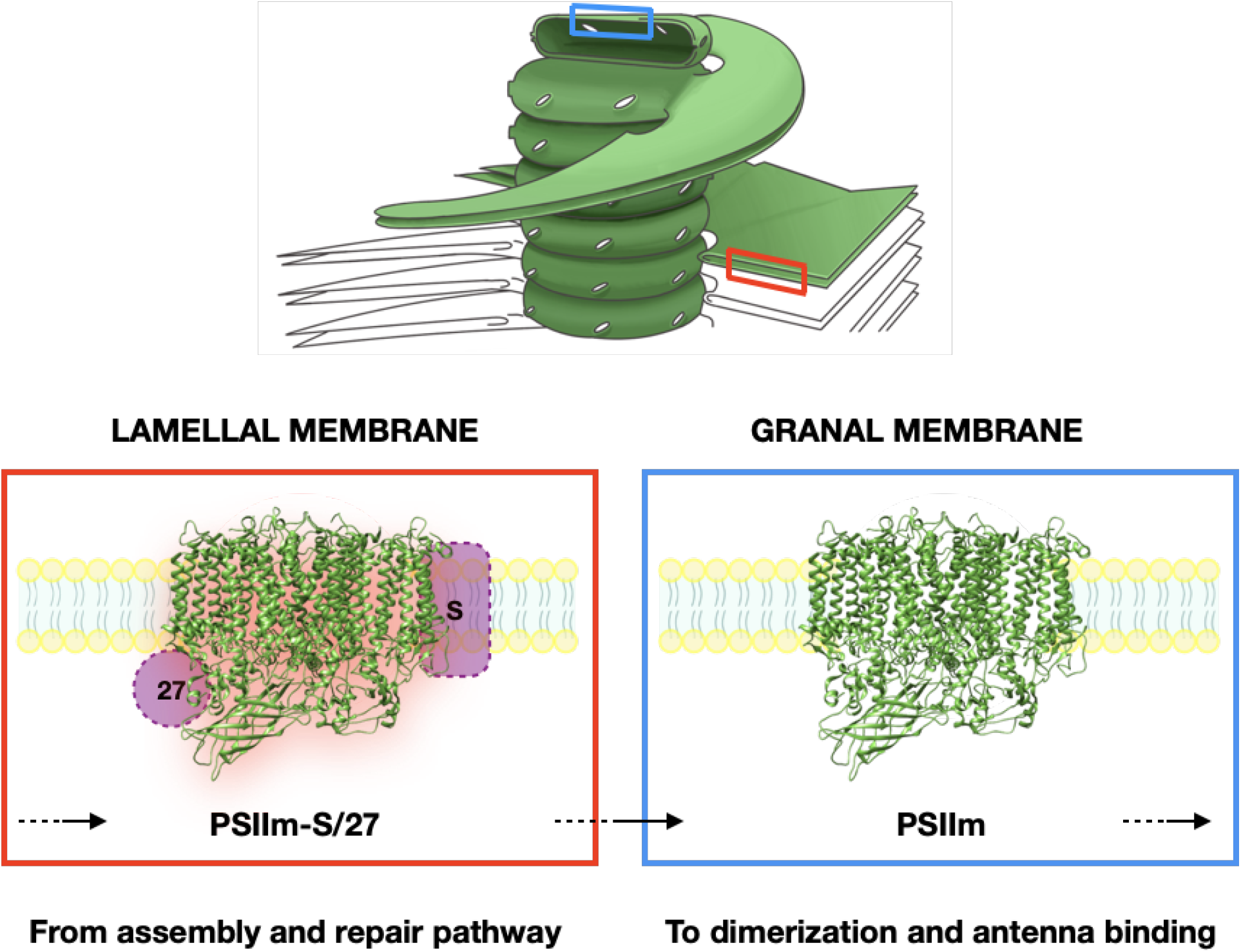
The location within the thylakoid membrane of the protein complexes investigated in this work. PSIIm-S/27 is in the lamella membranes (red inset) and suggested to be an intermediate in the assembly and repair pathway, probably after photoactivation. This complex migrates (as indicated by the black arrows) to the grana membranes (blue inset) where upon dissociation of PsbS and Psb27, it forms the active monomer, PSIIm. The PSIIm will then form active dimers and bind antenna proteins to form fully functional complexes.

PSII, the water/plastoquinone photo-oxidoreductase, uses the energy of light to drive charge separation, oxidise water and reduce plastoquinone. The photochemistry occurs as a one-photon/one-electron reaction four times sequentially to accumulate the 4 oxidising equivalents necessary for water oxidation and oxygen release at the Mn_4_CaO_5_ active site on the luminal side of PSII (Dau and Zaharieva, 2009). An exchangeable quinone, Q_B_, accepts 2 electrons and 2 protons sequentially, before it is released as Q_B_H_2_ from the stromal side of PSII into the PQ/PQH_2_ pool in the membrane (De Causmaecker et al., 2019). The sequential electron transfer steps involve the formation of a stable intermediate, Q_B_^•−^, that can back-react via Q_A_^•−^ with the semi-stable, charge accumulation intermediates of the Mn_4_CaO_5_ cluster (Rutherford et al., 1982). This back reaction occurs via the thermal repopulation of the P^•+^Pheo^•−^ state, which recombines mainly by a route forming the chlorophyll triplet state ^3^P_680_ (Rutherford et al., 1981). This triplet state reacts with oxygen to form singlet oxygen ^1^O_2_ (Krieger-Liszkay, 2005; Rutherford et al., 2012). The ^1^O_2_ generated causes damage to PSII (Krieger-Liszkay, 2005). Other reactive oxygen species, generated by reductive and oxidative processes in PSII, might also contribute to damage (Pospíšil, 2016).

Repairing the damage is an energetically costly process since proteins and cofactors must be synthesized and replaced to maintain efficient photosynthetic activity (Komenda et al., 2012; Tikkanen and Aro, 2012). This takes place in the stromal lamellae via a stepwise assembly of subcomplexes (Komenda et al., 2012; Nickelsen and Rengstl, 2013; Tomizioli et al., 2014; Puthiyaveetil et al., 2014). A large variety of PSII protein complexes are present in the stroma lamellae, with partially assembled systems co-existing with the PSII in different oligomeric states and different levels of activity (Danielsson et al., 2006; Haniewicz et al., 2013; Tomizioli et al., 2014; Puthiyaveetil et al., 2014). Inactive PSII from the stromal lamellae have been studied for decades (Melis, 1985; Lavergne and Leci 1993), and it was reported that 10-20% of PSII in the chloroplast were inactive and this was due to blocked forward electron transfer and not due to a lack of oxidised plastoquinone (Lavergne and Leci, 1993).

The complexity of the assembly/repair cycle, together with the low abundance of most of the intermediate complexes, means that our understanding of it is still evolving (Komenda et al., 2012; Nickelsen and Rengstl, 2013). Due to the low concentration, instability and intrinsically transient nature of these assembly intermediates, their isolation has required specific strategies: 1) the generation of mutants that lack either specific assembly factors or small PSII subunits, resulting in the accumulation of assembly intermediates (Komenda et al., 2002; Roose and Pakrasi, 2008; Zabert et al., 2021; Huang et al., 2021); and/or 2) tagging one of the PSII subunits to allow isolation of low concentration intermediates by affinity chromatography (Nowaczyk et al., 2006; Liu et al., 2011). Differential fractionation (Danielsson et al., 2006) and more recently differential solubilisation (Fey et al., 2008; Haniewicz et al., 2013) allowed the isolation of some of the sub-populations of PSII. Fractions originating from the lamellae and the granal margins (Haniewicz et al., 2013; Haniewicz et al., 2015) yielded a monomeric PSII containing two additional subunits, PsbS and Psb27, which are absent in functional granal PSII (Haniewicz et al., 2015). While Psb27 has been shown to bind to PSII sub-complexes and to play a role in PSII assembly (Nowaczky et al., 2006; Roose et al., 2008), PsbS has been associated, directly or indirectly, with photoprotection mechanisms via non-photochemical fluorescence quenching (Niyogi and Truong, 2013; Ruban et al., 2016; Bassi and Dall’Osto, 2021).

Since its discovery in 1984 (Ljungberg et al., 1984), the role of PsbS has been controversial (Niyogi and Truong, 2013; Fan et al., 2015; Ware et al., 2015; Ruban et al., 2016; Dall’Osto et al., 2017; Bassi and Dall’Osto, 2021). Despite the availability of a PsbS crystallographic structure (Fan et al., 2015), there is still a debate about its basic components. The presence of chlorophyll, xanthophyll (Correa-Galvis et al., 2016; Gachek et al., 2019) and its role as a luminal pH sensor (Bergantino et al., 2003; Li et al., 2004; Roach and Krieger-Liszkay, 2012; Liguori et al., 2019), are still debated. The primary role of PsbS is thought to be a protective one, as a key player in some aspects of non-photochemical quenching (NPQ) (Niyogi and Truong, 2013; Ruban et al., 2016; Bassi and Dall’Osto, 2021). Several reports showed the involvement, either direct or indirect, of PsbS, in quenching the excess of energy in free-LHCII complexes and/or in LHCIIs associated with PSII (Sacharz et al., 2017). A photo-protective role has also been suggested to act via CP47 and the minor external antennas of PSII (Correa-Galvis et al., 2016). However, to date, there is no consensus on a mechanism linking PsbS with NPQ and the xanthophyll cycle that would explain PsbS-mediated photoprotection and its role as a pH sensor.

PsbS has been found to be bound stoichiometrically to purified PSII cores only in samples originating from the lamellae and granal margins of *Tobacco* thylakoids (Haniewicz et al., 2013). Indirect evidence of its presence in PSII dimers and monomers, and its association to the PSII-LHCII complexes in grana have also been reported (Bergantino et al., 2003; Caffari et al., 2009; Correa-Galvis et al., 2016).

Psb27 is present in eukaryotes and prokaryotes, though most of the information relates to the cyanobacterial form. Important differences, such as the eukaryotic Psb27 lacking the covalently bound lipid moiety that is present in the cyanobacteria, raise doubts on whether they have the same location and function. In cyanobacteria, Psb27 is involved in the assembly of the Mn_4_CaO_5_ cluster and was found to be associated with inactive PSII lacking the three extrinsic proteins PsbO, PsbU and PsbV (Roose et al., 2004; Nowaczyk et al., 2006). It was found to play a significant role during PSII D1 repair, where it was suggested to bind to CP43 and facilitate the assembly of the Mn-cluster by providing greater accessibility and preventing premature association of the other extrinsic proteins (Nowaczyk et al., 2006; Roose and Pakrasi, 2008). Its location close to the CP43 loop E and its allosteric role in weakening the binding of the extrinsic proteins (Liu et al., 2011) was confirmed and clarified in two recent cryo-EM structures (Zabert et al., 2021; Huang et al., 2021). Its role in facilitating the photoactivation of the Mn-cluster was found to be more complex than simply displacing the extrinsic proteins from the apo-PSII (Avramov et al., 2020). In higher plants, this subunit is found to exist in two isoforms, Psb27-1 and Psb27-2 (Chen et al., 2006; Wei et al., 2010). Their specific function is still under investigation, with Psb27-1 found to be required for the efficient repair of photo-damaged PSII (Chen et al., 2006) but also to play a role in the state transition mechanism (Dietzel et al., 2011), while Psb27-2 is suggested to play an important role in the processing of the precursor form of D1 (Wei et al., 2010).

In the present study we have characterized a PSII monomer containing PsbS and Psb27, which was isolated from the stromal lamellae and the grana margins (PSIIm-S/27). We compared this complex with the PSII monomers isolated from the grana stacks (PSIIm). The data indicate an unexpected role for PsbS and/or the Psb27 protecting newly assembled and photoactivated PSII by inhibiting electron transfer, an inhibition that is reversed by bicarbonate binding.

## Results

### Association of PsbS and Psb27 to PSII monomers

Two types of purified Photosystem II core complexes, PSIIm and the PSIIm-S/27, were isolated according to the procedures previously described in Haniewicz *et al*. (2013) and in Haniewicz *et al*. (2015), where it was demonstrated that the PSIIm and the PSIIm-S/27 originate from the grana stacks and margins/stromal lamellae, respectively (Fig 1). The two types of monomeric PSII were compared by SDS-PAGE and we confirmed the previous observation that they had similar composition in terms of protein subunits, except for the two additional bands in the PSIIm-S/27 at 20 and 13 kDa (Fig. 2A), attributed to the PsbS and Psb27 subunits, respectively (Haniewicz *et al*., 2013). These subunits were found to be stoichiometric with the other core subunits of PSII. Small differences in the sub-stoichiometric content of external antennas, which are mainly ascribed to CP26 and CP29, were also observed (see SI Appendix).

**Figure 2:**
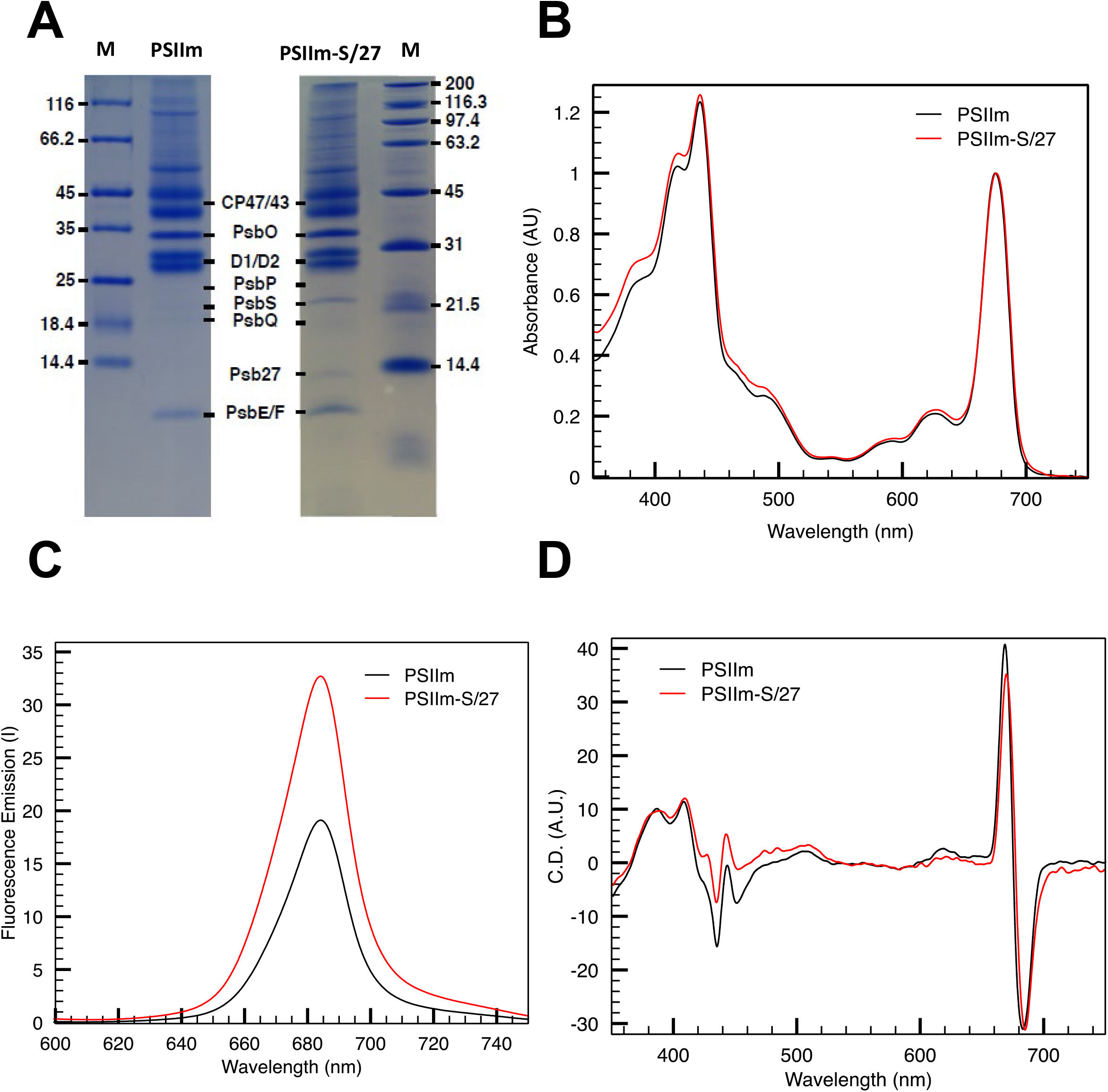
Comparisons of the PSIIm and PSIIm-S/27. A) Coomassie Blue Stained SDS-PAGE of the PSIIm (A) and PSIIm-S/27 (B). The additional presence of a ~ 22 kDa (PsbS) and a ~ 14 kDa (Psb27) bands in the PSIIm-S/27 sample are indicated together with the main PSII subunits. The lanes labelled as M indicate the molecular weight markers. B) UV-Vis absorption spectra of PSIIm (black line) and PSIIm-S/27 (red line). Spectra were taken at 20°C in buffer A and were normalized at 675 nm. C) Fluorescence emission spectra of PSIIm (black line) and PSIIm-S/27 (red line). Spectra, recorded at 4°C in buffer A with excitation at 437 nm. D) Circular dichroism spectra at 20°C in buffer A of PSIIm-S/27 (red line) and PSIIm (black line).

### Spectroscopic characterisation of PSIIm and the PSIIm-S/27

UV-Vis absorption spectra of the PSIIm and the PSIIm-S/27 were recorded (Fig. 2B). When normalising the spectra at 675 nm, the comparison showed only minimal differences localised between 350 and 550 nm with the PSIIm-S/27 showing slightly higher absorbance. As neither the Psb27 and PsbS have been shown to contain chromophores, these differences are more likely to be due to the small differences in CP26 and CP29 content of the samples.

Room-temperature fluorescence emission spectra were recorded for both PSII samples and showed a single peak at 681 and 682.5 nm for PSIIm-S/27 and PSIIm, respectively (Fig. 2C). In equally concentrated samples, the intensity of the emission peak was much higher (nearly double) for the PSIIm-S/27 sample when compared with the PSIIm (Fig. 2C).

The comparison of the circular dichroism spectra (Fig. 2D) for the PSIIm and the PSIIm-S/27 samples showed differences that can be related to the small changes in the absorption spectra shown in the Fig. 2B. Overall, both spectra resemble a typical PSII spectrum (Alfonso et al., 1994; Kraus et al., 2005), suggesting that no major changes in the position or number of the cofactors is induced by the presence of PsbS and Psb27. However, in the PSIIm-S/27 sample the spectral region between 350 and 550 nm showed changes in peak intensity and position, with 3 minor bands at 427, 475, and 484 nm, respectively. The Qy band showed the typical PSII double peak in both samples (Alfonso et al., 1994; Kraus et al., 2005). In the PSIIm-S/27 sample, these features were red-shifted by about 2 nm with respect to PSIIm (Fig. 2D).

### Water oxidation catalytic activity

The enzymatic activities of both PSIIm and PSIIm-S/27 were compared by measuring their oxygen evolution rates. PSIIm showed good activity with rates of 1030±13 μmol O_2_ mgChl^-1^ h^-1^. In contrast, PSIIm-S/27 showed a drastically reduced activity of 52±5 μmol O_2_ mgChl^-1^ h^-1^ (Table 1).

**Table 1:** Rates of oxygen evolution for PSIIm and PSIIm-S/27 with or without added bicarbonate. Data represent mean +/- SD, n = 3.

| PSII type | Added [NaHCO<sub>3</sub>] | Oxygen evolution rates<br>($\mu\text{mol O}_2/\text{mg Chls}\cdot\text{h}$) |
| --- | --- | --- |
| PSII<sub>m</sub> | 0 | 1030 ± 13 |
| PSII<sub>m</sub>-S/27 | 0 | 52 ± 5 |
| | 5 mM | 795 ± 8 |

### Electron transfer from Q_A_^•−^ to Q_B_ or Q_B_^•−^

PSII photochemistry was tested by measuring flash-induced chlorophyll fluorescence and Q_A_^•−^ oxidation kinetics. The illumination with a single-saturating flash of a dark-adapted sample induces the reduction of Q_A_ to Q_A_^•−^ in most of the centres, with a resulting increase in the prompt fluorescence yield. Subsequent re-oxidation of Q_A_^•−^, either by electron transfer to Q_B_ or Q_B_^•−^ or by recombination with S_2_, results in the decay of the fluorescence yield (Croft and Wraight, 1983).

Figure 3A shows that both minimal fluorescence (F_0_) and maximal fluorescence (F_m_) yields were higher in the PSIIm-S/27 sample (see SI Appendix). PSIIm was found to be unstable during the measurements at room temperature, therefore experiments were performed at 15°C (Fig. 3B and D). This is likely to slow down some rates when compared to other measurements done at room temperature and when comparing them to the literature. The decay rates of the fluorescence yield are shown in Fig. 3B. PSIIm kinetics are like those expected for functional PSII, while in PSIIm-S/27 the fluorescence decayed ~10 times more slowly, indicating a marked inhibition of forward electron transfer. The kinetics were fitted with three decay phases (Fig. 3B and Table 2) and the origins of the decay phases were assigned to forward and backward electron transfer reactions according to the literature (Vass et al., 1999).

**Figure 3:**
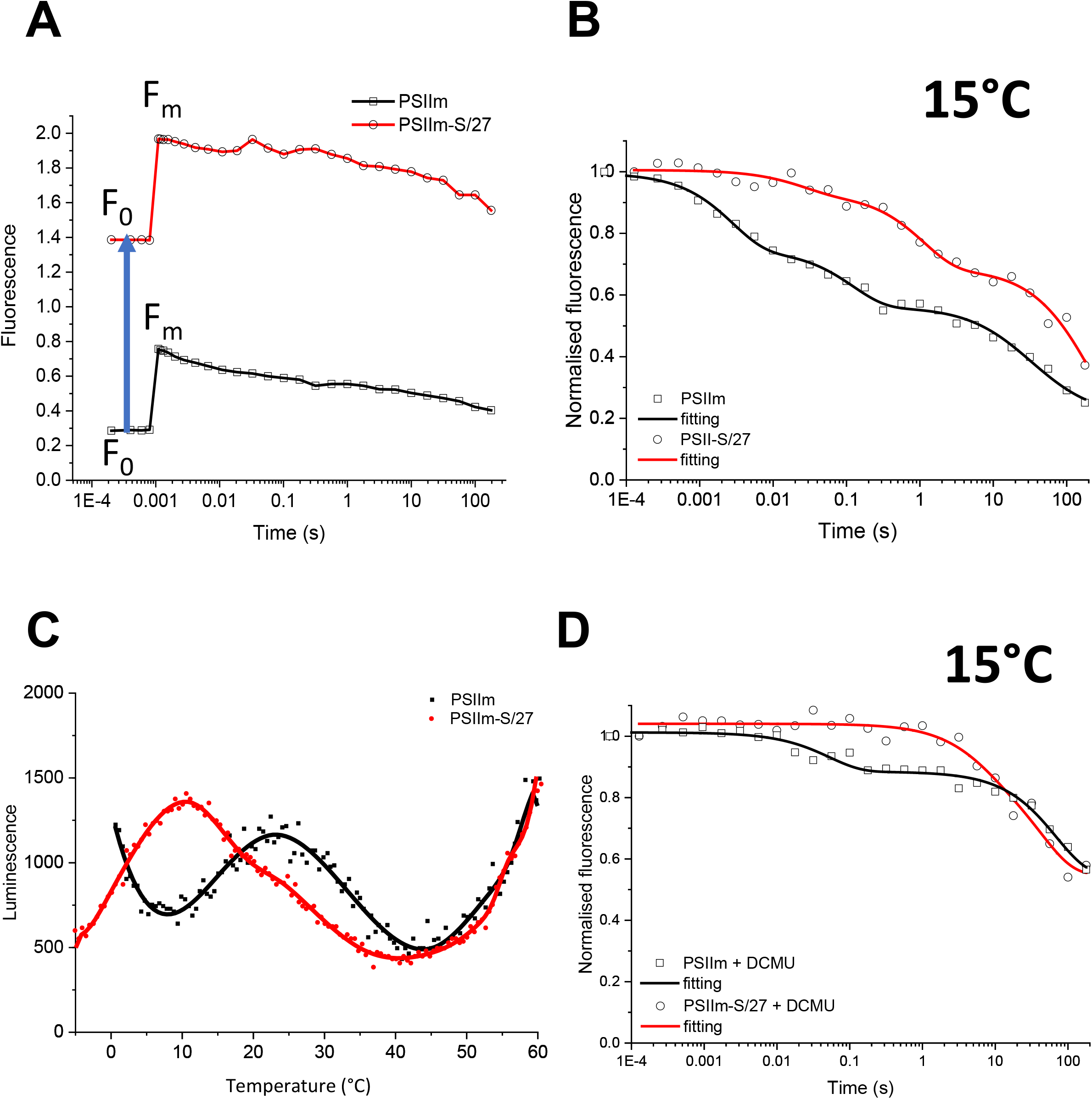
A) Fluorescence relaxation kinetics data presented without normalization to show the values of F_0_ and F_m_ for PSIIm (squares symbols, black line) and PSIIm-S/27 (circles symbols, red line). Both F_0_ and F_m_ were found to be higher in the sample with bound PsbS and Psb27. B) Fluorescence relaxation kinetics of PSIIm (squares, black line) and PSIIm-S/27 (circles, red line), measured at 15°C in buffer A. Data were normalized using the initial amplitudes. Fittings were carried out with equation 1 (see methods). C) Thermoluminescence measurements for PSIIm (black squares and line) and PSIIm-S/27 (red circles and line), in buffer A. Single saturating flash was given at 5°C followed by rapid cooling to −5°C. Scan rate was 0.5°C/s. D) Fluorescence relaxation kinetics upon a single saturating flash for the PSIIm (squares, black line) and PSIIm-S/27 (circles, red line), measured at 15°C in buffer A in the presence of 10 μM DCMU. Data were normalized using the initial amplitudes. Fittings were carried out with equation 1 (see methods).

**Table 2:**
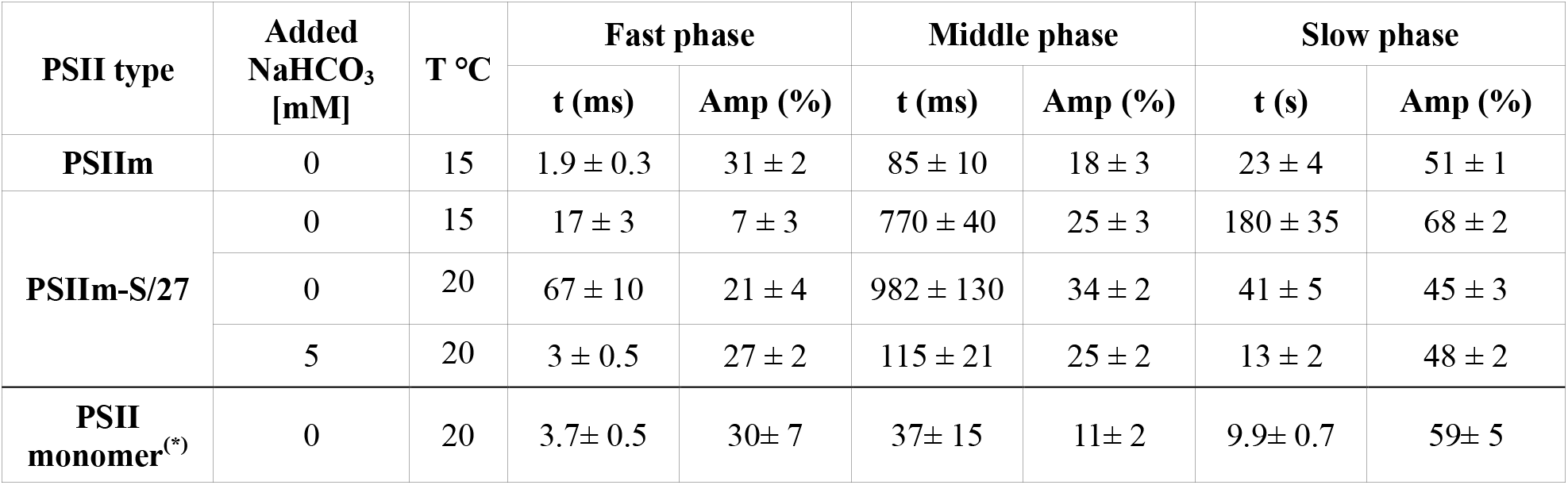
Kinetic parameters of flash-induced chlorophyll fluorescence decay in PSIIm, PSIIm-S/27 samples in absence or presence of additional bicarbonate in solution. The values of the kinetic half-lives (t) and the amplitudes of each phase are compared to the literature values for PSII monomer from *T. elongatus* (Zimmermann et al., 2006)(*). Data represent mean +/- SD, n = 3.

The initial fast phase arises from Q_A_^•−^ forward electron transfer to either Q_B_ or Q_B_^•−^. It has a rate of 1.9 ms and an amplitude of 31% for the PSIIm, but a drastically reduced rate and amplitude of 17 ms and 7% amplitude for PSIIm-S/27.

The middle phase is often assigned to be electron transfer from Q_A_^•−^ when the Q_B_ site is either empty or occupied by Q_B_H_2_ at the time of the flash, and therefore the electron transfer rate is determined by the arrival of PQ into the Q_B_ site. For PSIIm this phase shows an amplitude of 18% and a half-time of 85 ms, kinetics compatible with the usual assignment of this phase although on the slow side of the range and could indicate a contribution to this phase of charge recombination with TyrZ• in damaged centres. However, for PSIIm-S/27, while the amplitude is like that in PSIIm, the decay is 9-fold slower, with a t_1/2_ = 770 ms. This is a very slow value for quinone exchange although slow quinone exchange is a feature of bicarbonate loss from the non-heme iron or bicarbonate replacement by other carboxylic acids (Shevela et al., 2012). This range of fluorescence decay is within those seen for S_2_ recombination with Q_A_^•−^ in PSII monomers (Zimmermann et al., 2006) but it seems that in this material and at this temperature, this S_2_Q_A_^•−^ recombination takes place more slowly (see below).

The third and slowest phase is usually attributed to the back reaction of Q_A_^•−^ with the S_2_ state of the Mn-cluster and thus this phase is seen when forward electron transfer is blocked due to reduction of the pool, modification of the Q_B_ site or binding of an herbicide in the Q_B_ site. This phase with a t_1/2_ = 23s seems to be present in about half of the centres in PSIIm, this is in common with other reports from PSIIm in the literature (Table 2). The slower rates compared with the literature values is due to the lower temperature (15°C) used here. The PSIIm-S/27 sample showed even more of this phase, 68%, and a marked slow-down of the half-time to 180s. Thermoluminescence measurements (TL) of PSIIm showed that upon a single saturating flash the PSIIm presents a single peak at 22°C, while PSIIm-S/27 presents a peak at 10°C with a shoulder at 25°C (Fig. 3C and S1). These peaks may be attributed to S_2_Q_A_^•−^ and S_2_Q_B_^•−^ recombination respectively based on the typical TL peak temperatures (Rutherford et al 1982). The TL data suggest that the presence of PsbS and Psb27 block the electron transfer from Q_A_^•−^ to Q_B_. This appears to agree with fluorescence kinetics presented above on the Q_A_^•−^ re-oxidation kinetic considering the temperature of the measurement.

### Effect of the inhibitor DCMU on the Q_A_^•−^ oxidation kinetics

To investigate the possible interference of PsbS and/or Psb27 with Q_B_ binding, we measured the fluorescence yield relaxation kinetics in the presence of the PSII inhibitor 3-(3,4-dichlorophenyl)-1,1-dimethylurea (DCMU) (Fig. 3D). This inhibitor binds to the Q_B_ binding site and blocks the electron transfer from Q_A_^•−^, leaving the recombination to S_2_ as the only possible route for the electrons. The kinetics of Q_A_^•−^ oxidation will, therefore, be dominated by the slow phase associated with the recombination with S_2_. Addition of DCMU to both the PSIIm and the PSIIm-S/27 resulted in kinetics with very similar half-times of approximately 30-40 s in 50-60% of the centres, while the remaining 40-50% appear to show longer decaying times. This observation suggests that the presence of PsbS and/or Psb27 does not interfere with the DCMU binding nor the resulting inhibition. These rates are longer than those typically measured in fully functional plant PSII, where it is ~1 second, but this is at least partially explained by the experiments being done at 15°C to preserve the intactness of PSIIm. The slow phases of S_2_Q_A_^-^ decay measured with DCMU are similar in PSIIm and PSIIm-S/27, while in the absence of DCMU the slow phase of fluorescence decay was significantly slower in PSIIm-S/27 (Fig. 3B and D).

A notable difference in the kinetics is shown for the PSIIm in the presence of DCMU, where an additional faster phase is present (Fig. 3D) with ~20% amplitude and t_1/2_ = 50 ms. This phase could correspond to Tyr_z_•Q_A_^•−^ recombination, a reaction reflecting PSII centres that lack the Mn-cluster. This is consistent with the observed instability of the PSIIm.

### Effect of bicarbonate on PSIIm-S/27

Given the recently discovered protective role of bicarbonate and the demonstration that it can be lost under physiological conditions (Brinkert et al., 2016), we investigated the effect of bicarbonate on PSIIm-S/27. These experiments were done at 20°C, a temperature at which the effect of bicarbonate has been characterised and the PSIIm-S/27 was stable. The addition of 5 mM bicarbonate to the PSIIm-S/27 sample resulted in it becoming activated to a level comparable to the functional PSIIm (Fig. 4A). The kinetics of Q_A_^•−^ oxidation after addition of bicarbonate showed an acceleration of all three fluorescence decay phases to rates like those measured in the PSIIm and typical of a functional PSII monomer (Zimmermann et al., 2006). In the presence of bicarbonate, the fast phase t_1/2_ decreased from 67 ms to 3 ms, the middle phase decreased from 982 ms to 115 ms, and the slow phase from 43 to 13 s (Table 2). The amplitude of the fast phase also appeared to increase when bicarbonate was present, but this is less certain because of the influence of the poorer fitting of the equivalent phase but with slower kinetic, in the PSIIm-S/27 lacking bicarbonate.

**Figure 4:**
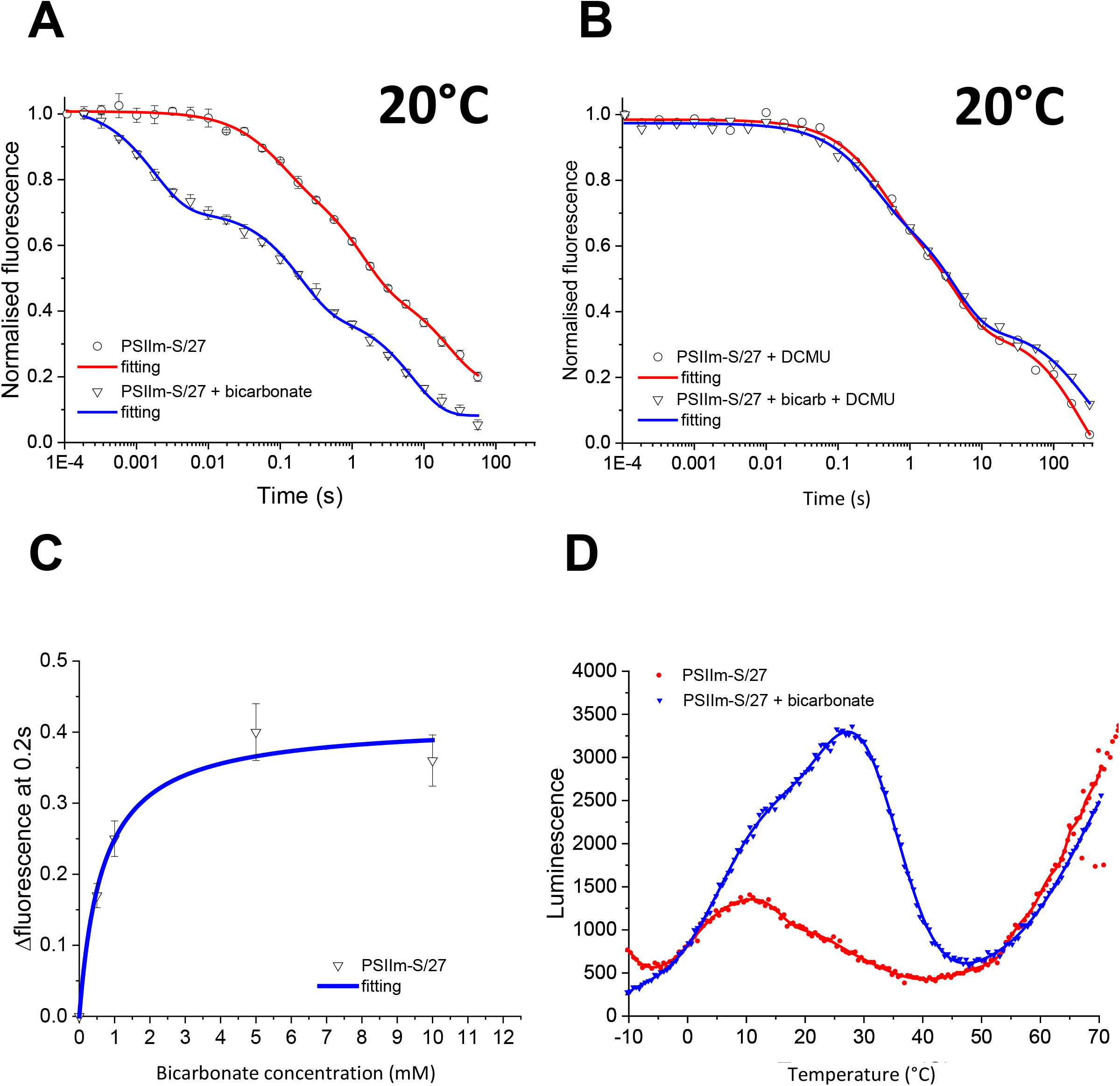
A) Fluorescence relaxation kinetics for PSIIm-S/27, measured at 20 °C in buffer A, without (circles symbols, red line) and with (triangles symbols, blue line) added 5 mM bicarbonate. Data were normalized using the initial amplitudes. Fittings were carried out with equation 1 (see methods). Error bars represent the standard errors calculated from 4 independent measurements. B) Fluorescence relaxation kinetics for PSIIm-S/27, measured at 20 °C in buffer A and 10 μM DCMU, without (circles symbols, red line) and with (triangles symbols, blue line) added 5 mM bicarbonate. Data were normalized using the initial amplitudes. Fittings were carried out with equation 1 (see methods). Error bars represent the standard errors calculated from 4 independent measurements. C) Plot of the fluorescence intensity at 0.2 s (triangles symbols) from the fluorescence relaxation kinetics, at 20 °C in buffer A, from different samples to which increasing concentrations of bicarbonate were added. The data were fitted with the hyperbole in equation 2 (see methods) (blue line). Error bars represent the standard errors calculated from 4 independent measurements; D) Thermoluminescence measurements of PSIIm-S/27 in buffer A, in the absence (red circles) and presence (blue triangles) of added 5 mM bicarbonate. Single saturating flash was given at 5 °C followed by rapid cooling to −5 °C. Scan rate was 0.5 °C/s.

The addition of DCMU to the bicarbonate containing PSIIm-S/27 complex yielded almost the same kinetic profile as seen in the absence of the added 5 mM bicarbonate (Fig. 4B). The fitting of the data showed two main phases, the first with a half-time of approximately 1 s and an amplitude of 70%, consistent with S_2_Q_A_^•−^ recombination, the second with a half-time of approximately 60 s and an amplitude of 30%. The longer half-life of this second phase is consistent with the oxidation of a relatively stable donor (e.g. from Mn^2+^, TyrD, the side path donors) giving rise to a long-lived Q_A_^•−^ (Nixon et al., 1992; Debus et al., 2000).

Oxygen evolution assays performed in presence of 5 mM bicarbonate showed a ~15-fold increase in the activity of PSIIm-S/27, reaching levels (795±8 μmol O_2_ mg Chl^-1^ h^-1^) that are ~70% of the values recorded for the PSIIm (Table 1). These results show that the PSIIm-S/27 samples contain a fully functional Mn-cluster in at least 70% of the centers. Both kinetics and oxygen evolution measurements show that approximately 30% of the centers lack catalytic activity and these show a kinetic profile that is consistent with either partial Mn occupancy or Mn-free PSII.

The kinetics of oxidation was studied as a function of bicarbonate concentration. The kinetics accelerated as the bicarbonate concentration was increased. When the fluorescence value at 0.2 s were plotted, and the value obtained prior to bicarbonate addition was subtracted (Fig. 4C), the curve showed hyperbolic behavior saturating at 5 mM. Data fitting with a hyperbolic model for ligand binding yielded an apparent dissociation constant for the bicarbonate of ~ 600 μM.

TL (Fig. 4D) of PSIIm-S/27 with no addition showed a peak centered at 10°C attributed to S_2_Q_A_^•−^ recombination, while in the presence of bicarbonate the TL intensity increased with a dominant peak at 27°C, typical of S_2_Q_B_^•−^ recombination, and an increase in S_2_Q_A_^•−^ TL at 10°C, which is seen as shoulder. The TL results show that PSIIm-S/27 has inhibited forward electron transfer from Q_A_^•−^ and only a low level of luminescence arising from S_2_Q_A_^•−^. Addition of bicarbonate resulted in the recovery of near-normal behavior with the formation of S_2_Q_B_^•−^ recombination in most of the centres. In a fraction of centres the bicarbonate did not reconstitute electron transfer to Q_B_^•−^ but did result in more S_2_Q_A_^•−^ recombination.

Finally, it was also observed that the peak intensity and position of the room temperature fluorescence was essentially unaffected by the addition of 5 mM bicarbonate (Fig. S2).

## Discussion

Here, we compared two types of PSII monomers, i) those isolated from grana, PSIIm, and ii) those isolated from the stromal lamellae and granal margins, which contain stoichiometric amounts of PsbS and Psb27, PSIIm-S/27 (Haniewicz *et al*., 2013; Haniewicz *et al*., 2015). Comparison of the UV-Vis absorption and CD spectra (Figs. 2B and 2D) in the two types of PSII monomers, showed only minimal differences (see SI Appendix). The most significant difference between the two complexes is the near absence of activity in PSIIm-S/27 compared to the high activity in the PSIIm. Both the oxygen evolution rates (Table 1) and the kinetics of Q_A_^•−^ oxidation (Fig. 3 and Table 2) were strongly inhibited in PSIIm-S/27. The addition of DCMU to PSIIm-S/27 blocked oxygen evolution and shut down the residual, sub-second fluorescence decay, due the near-complete block of forward electron transfer from Q_A_^•−^. The DCMU-treated PSIIm-S/27 showed 70% of the centres with the typical seconds-timescale kinetics of S_2_Q_A_^•−^ recombination (Fig. 3D and 4B), while the rest of the centres showed much slower rates of Q_A_^•−^ decay, presumably due to the electron donation from a more stable electron donor in a fraction of centres. These observations are comparable to those made by Lavergne and Leci (1993) when investigating the fraction of inactive PSII that is normally present in green algal cells. The inactive PSII in algal cells reported earlier (Lavergne and Leci, 1993) could be the algal equivalent of the PSIIm-S/27 described here.

The kinetic characteristics of the impaired Q_A_^•−^ oxidation in PSIIm-S/27 indicate heterogeneity and suggest that forward electron transfer, Q_A_^•−^ to Q_B_ and to Q_B_^•−^, and the exchange of Q_B_H_2_, are all inhibited. These events all involve protonation. The thermoluminescence of PSIIm-S/27 had a low intensity but the peak positions of the residual TL were consistent with inhibition of electron transfer from Q_A_^•−^ to Q_B_ or Q_B_^•−^, with the main peak at 10°C, typical of S_2_Q_A_^•−^ recombination in inhibited centres, and only a very small shoulder at 25°C corresponding to the S_2_Q_B_^•−^ recombination in a small number of functional centres (Fig. 3C).

The inhibition of Q_A_^•−^ oxidation in PSIIm-S/27 cannot be explained by the loss of Q_B_ (except in a small fraction of the centres), as the addition of bicarbonate activated forward electron transfer in most of the centres. Similarly, the weak TL intensity cannot be explained by the absence of the Mn cluster, as most of the centres were capable of water splitting when bicarbonate was added. The low luminescence of PSIIm-S/27 prior to the addition of bicarbonate, and despite the presence of the Mn cluster and both quinones, could be due to an increase in the redox potential Q_A_^•−^, as occurs upon loss of the bicarbonate from granal PSII dimers (Brinkert et al., 2016). A sufficiently high redox potential is expected to result in the loss of radiative recombination (Rutherford et al., 2012).

The remarkable activation of the seemingly inactive PSIIm-S/27 by millimolar concentrations of bicarbonate was manifest as the appearance of normal rates of forward electron transfer and water oxidation activity in at least 70% of the centres, as monitored by fluorescence kinetics, TL and O_2_ evolution. The remaining 30% inactive centres did show a slow-down of the rate of Q_A_^•−^ reoxidation in the ~10 s range (Fig. 4A, Table 2). As these kinetics are much slower than typically found for electron transfer to Q_B_ and Q_B_^•−^, this observation could indicate the absence of Q_B_ in the site in this fraction of the centers. However, as the 10 s phase is eliminated in the presence of DCMU and replaced by a much slower rate, ~100 s (Fig. 4B), this behaviour could indicate an unusually slow rate of electron transfer from Q_A_^•−^ to Q_B_. This could originate from a situation where the reduction potentials of Q_A_/Q_A_^•−^ and Q_B_/Q_B_^•−^ are similar. The reduction potential of Q_A_/Q_A_^•−^ is reported to shift toward that of Q_B_/Q_B_^•−^ when the Mn cluster is absent (Johnson et al., 1995), and it could be shifted even further when modified by the binding of the PsbS and Psb27 subunits.

The major difference between the two types of PSII monomers studied here is that the PSIIm-S/27 almost completely lacks activity until activated by the addition of bicarbonate. This difference is presumably due to the binding of PsbS and/or Psb27. It is not clear that this difference is due to one or the other of these polypeptides or to a combination of both. This uncertainty is shared with the recent structural work on other assembly intermediates in cyanobacterial systems (Huang et al., 2021; Zebert et al., 2021). Below we discuss the potential roles of the Psb27 and PsbS.

Psb27 in higher plants is relatively poorly studied, and is known to exist in two isoforms, Psb27-1 and Psb27-2 (Chen et al., 2006; Wei et al., 2010). In the present study, it was not possible to determine which of the two isoforms is bound to PSIIm-S/27. Its suggested functions are associated with responses to photodamage and maturation of D1 in newly synthesised PSII (Chen et al., 2006; Wei et al., 2010). In cyanobacteria, Psb27 is involved in the assembly of the Mn_4_CaO_5_ cluster, where it is suggested to facilitate photoassembly by allosterically regulating the binding of the extrinsic proteins, PsbO, PsbU, and PsbV (Nowaczyk et al., 2006; Roose and Pakrasi, 2008). In both *Synechocystis* sp PCC 6803 and *Thermosynechococcus elongatus*, Psb27 was found to be associated with inactive PSII monomers and dimers in which the three extrinsic proteins were absent and either no Mn or sub-stoichiometric amounts of Mn were reported (Roose and Pakrasi, 2004; Nowaczyk et al., 2006; Mamedov et al., 2007; Roose and Pakrasi, 2008). Nevertheless, PSII complexes with Psb27 bound in the presence of either PsbO alone, or the full complement of extrinsic proteins, were found in a range of conditions: 1) as PSII dimers in cold-stressed *T. elongatus* (Grasse et al., 2011), 2) in affinity purified His-tagged Psb27 in *Synechocystis* sp PCC 6803 (Liu et al., 2011), and 3) as PSII monomers in a *psbJ* deletion mutant of *T*. elongatus (Zabert et al., 2021).

Two recent structures of PSII complexes with bound Psb27 (Zabert et al., 2021; Huang et al., 2021), confirmed the previously suggested (Liu et al., 2011; Komenda et al 2012) binding site for Psb27 close to the loop E in CP43. The structures also indicate that binding of Psb27 does not directly interfere with the binding of the extrinsic proteins, in agreement with the range of different Psb27-bound forms of PSII reported in the literature, and therefore reflecting the dynamic process of assembly and repair (Komenda et al., 2002; Roose and Pakrasi, 2004; Nowaczyk et al., 2006; Mamedov et al., 2007; Roose and Pakrasi, 2008; Grasse et al., 2011; Liu et al., 2011; Komenda et al., 2012; Avramov et al., 2020; Huang et al., 2021; Zabert et al., 2021). This agrees with the observation in the present work that in the PSIIm-S/27, the Psb27 is bound, the extrinsic polypeptides are also bound, and the Mn_4_CaO_5_ cluster is fully assembled in the majority (70%) of the centres. It is not clear if the PSIIm-S/27 is an early-stage intermediate in the repair/assembly process, as suggested in Grasse *et al*. (2011) for a Mn-containing but inactive Psb27-bound PSII dimer in *T.elongatus* (Grasse et al., 2011), or a late stage intermediate, following photoassembly of Mn cluster, prior to joining the fully functional PSII population in the grana. However, the high activity seen in PSIIm-S/27 when the centres were activated by bicarbonate, points to a lack of photodamage and favours its assignment as a late stage post-photoactivation intermediate.

The recent cryo-EM structure of the PSII monomer from *T. elongatus*, showing Psb27 bound to the CP43 and no Mn cluster, also showed significant modifications to the structure around the non-heme iron and the Q_B_ site. The acceptor side modifications appear to be related to the binding of two other polypeptides, the Psb28 and the Psb34, that cause a conformational change of the D-E loop of the D1 protein that forms a stabilising interaction with Psb28. Part of the C-terminus of CP47 is also displaced by Psb34 forming a stabilising interaction with Psb28. Perhaps the most remarkable result of this conformational change was that the bicarbonate ligand to the non-heme iron was displaced by the carboxylic group of Glu241-D2 (Xiao et al., 2021; Zabert et al., 2021).

Kinetics of Q_A_^•−^ oxidation measured for this complex (Zabert et al., 2021), and in other related complexes (Liu et al., 2011; Mamedov et al., 2011), all show large fractions of centres with slow Q_A_^•−^ decay (greater than 10 s). This slow decay of Q_A_^•−^ has been attributed to a situation in which forward electron transfer is blocked and Q_A_^•−^ is trapped due to a stable electron donor to either Tyr_z_• or P_D1_^+^, such as a Mn^2+^ or a side-path donor (Nixon et al., 1992; Debus et al., 2000). This situation resembles that observed here in the fraction (~30%) of centres of PSIIm-S/27 lacking the intact Mn cluster. This fraction could represent either centres that have yet to undergo photoactivation, similarly to the intermediates presented in Zabert *et al*. (2021), or centres that have lost the Mn-cluster during isolation from the thylakoid membrane.

Given the clear association of the Psb27 with the electron donor side in the cyanobacterial system, it is tempting to suggest that the plant Psb27 binds in a similar location and plays a similar role/s. It has been known for decades that changes on the electron donor side can have major effects on the electron acceptor side (Johnson et al., 1995). It is thus possible that the binding of the Psb27 has a long-range effect on the electron acceptor side. Indeed, in the crystal structure of the PSII dimer with Psb27, the Psb27 binding site overlaps with the binding site of PsbQ in plants, and PsbQ’ in red algae and diatoms (Huang et al., 2021). It has been reported that the PsbQ’ binding to PSII shifts the reduction potential of the QA/Q_A_^•−^ couple to more positive values (Yamada et al., 2018). This suggests the possibility that the binding of Psb27 might result in a shift in the reduction potential of Q_A_/Q_A_^•−^.

There is much less relevant information for the PsbS, as there are no examples of isolated PSII cores with bound PsbS other than the PSII monomer studied here (Haniewicz *et al*., 2013; Haniewicz *et al*., 2015). Previous work on PsbS has been aimed at understanding its role, directly or indirectly associated with non-photochemical quenching (Niyogi and Truong, 2013; Ruban, 2016; Bassi and Dall’Osto, 2021). However, the PSIIm-S/27 was isolated from plants that had not been subjected to light stress leading to the induction of non-photochemical quenching. Furthermore, the spectroscopic characterisation of PSIIm-S/27 (Fig. 2 and 3) shows no quenching of the fluorescence due to PsbS binding. Therefore, a different functional role for PsbS in the PSIIm-S/27 complex should be considered. Cross-linking experiments in thylakoid membranes upon induction of quenching, indicated that monomeric PsbS was associated with CP47 and D2, in addition to its expected association with LHCII (Correa-Galvis et al., 2016). The N-terminal and C-terminal loops, which are both on the stromal side of PsbS, could interact with the electron acceptor side of PSII and affect its function. In the absence of further structural information, it remains possible that the PsbS, in PSIIm-S/27 is located as suggested in the cross-linking experiments (Correa-Galvis et al., 2016).

Now we turn to the effect of bicarbonate. Here, we found that the addition of bicarbonate activated PSIIm-S/27 giving normal rates of Q_A_^•−^ oxidation (Fig. 4A and Table 2) and oxygen evolution (Table 1) and near normal TL (4D, Fig. S2). This unexpected bicarbonate-dependent activation showed that the PSIIm-S/27 complex as isolated was essentially intact and capable of normal function but was “switched off”. The study of the kinetics of Q_A_^•−^ oxidation as a function of the bicarbonate concentration showed a hyperbolic dependence, typical of the binding of a ligand to a discrete binding site and an apparent dissociation constant of ~ 600 μM. Brinkert *et al*. (2016) showed that the bicarbonate binding site on the non-heme Fe in functional granal PSII dimers has two binding affinities: a high affinity when Q_A_ is present and a lower affinity when long-lived Q_A_^•−^ is present. Based on the thermodynamic relationship between the effect of bicarbonate binding and the reduction potential (E_m_) of Q_A_/Q_A_^•−^, and the literature *Kd* of 80 μM (Stemler and Murphy, 1983) taken as the high affinity value, Brinkert *et al*. (2016) calculated the *Kd* for the low affinity state to be 1.4 mM. However, they pointed out that based on their observations, the literature *Kd* value, 80 μM, appeared to be overestimated and suggested that the actual value was in the low μM range, i.e., that it had a significantly higher affinity. This would mean that the *Kd* for the low affinity conformation would be smaller than the 1.4 mM, and therefore close to, or smaller than the 600 μM *Kd* measured here for bicarbonate activation of PSIIm-S/27.

Brinkert *et al*. (2016) argued that the increase in the E_m_ of Q_A_/Q_A_^•−^ that occurred upon the loss of the bicarbonate, would increase the energy gap between Pheo and Q_A_, and this would disfavour the back-reaction route for charge recombination and favour direct charge recombination (Johnson et al, 1995). As described in the introduction, the back-reaction route via the pheophytin leads to chlorophyll triplet formation and thence to ^1^O_2_-mediated photodamage, while the direct recombination of the slower P^+^Q_A_^•−^ radical pair is considered safe (Rutherford et al., 1982; Johnson et al., 1995). The bicarbonate-mediated redox tuning of Q_A_ was thus considered to be a regulatory and protective mechanism (Brinkert et al., 2016). It seems quite likely that a similar protective mechanism exists in the PSIIm-S/27, as protection is needed during and after synthesis or repair.

The increased stability of the PSIIm-S/27 conferred by the presence of PsbS and/or Psb27 was manifest by its resilience to long incubations and photochemical measurements at room temperature. This resilience contrasted with the fragility of the PSIIm, which could not be studied even for short periods at room temperature without loss of activity.

A bicarbonate-controlled, redox-tuning-based, protective mechanism in PSIIm-S/27 would appear to be beneficial for this complex. A nonfunctional PSII, like PSIIm-S/27 with an intact electron donor-side but with inhibited forward electron transfer and a low-potential Q_A_ (Johnson et al., 1995), would be hypersensitive to backreaction-associated photodamage, just as occurs in herbicide-treated PSII (Rutherford and Krieger-Liszkay, 2001). It is known that before photoactivation of water oxidation in PSII, the E_m_ of Q_A_/Q_A_^•−^ is high and thus PSII is photo-protected, and at some point during the photoassembly of the Mn complex, the E_m_ is shifted to a functional, low potential value (Johnson et al., 1995). If the PSIIm-S/27 is a late-stage intermediate of photoactivation, then the present work would indicate that the donor-side-induced switching of the E_m_ of Q_A_/Q_A_^•−^ is overridden by modifications to the acceptor side that maintain Q_A_ in a safe, high-potential form, until the fully assembled PSII is delivered into the granal stack and is dimerised. The binding of the PsbS and Psb27, can be considered as exerting conformation restrictions to the assembled but non-functional PSII, protecting it until it is in the right place and dimerised. At that point, presumably the PsbS and Psb27 dissociate, allowing the PSII to adopt its functional conformation, allowing the high affinity bicarbonate site to form. The bicarbonate duly binds, shifting the E_m_ of Q_A_/Q_A_^•−^ to low potential and allowing optimal function. When considering the measured dissociation constant for bicarbonate in PSIIm-S/27 (600 μM), it seems clear that the physiological concentration of CO_2_ could control, to some extent, the activity of this complex. At pH 8.0 the equilibrium concentration of bicarbonate will be sufficiently high to allow more than 50% of this complex to show normal forward electron transfer kinetics.

Another less likely explanation for the lower activity of PSIIm-S/27 is that it represents an early-stage intermediate in the repair cycle. In this model, photodamage would be manifest as an electron acceptor side restriction, and the binding of Psb27 and PsbS and the higher potential form of Q_A_, due to the loss of bicarbonate, would protect the system from photodamage during its transit to the repair site in the stromal lamellae.

In the literature there are several examples of other PSII subunits which seem to exhibit similar or related effects to those described here for PsbS and Psb27, though none of them are as marked as observed in this work. These are listed in the supplementary information (See SI Appendix). This evidence in the literature and the present work point to a broader picture in which there is an interplay between the binding of small subunits to PSII with the resulting conformational effects, and the binding of bicarbonate with the resulting redox tuning effects. This interplay appears to control electron transfer rates and thermodynamic equilibria between the different quinones in all their forms, thereby regulating and safeguarding PSII during the diverse steps of its life cycle.

## Conclusions

We show that the PSII monomer from the stromal lamellae/stromal margins, which has PsbS and Psb27 bound to it, has very low activity but is activated upon binding bicarbonate. These findings indicate that PSIIm-S/27 is a switched-off state that is protected from photodamage presumably due to the changes induced by the binding of the two extra polypetides. The nature of the protection mechanism appears to be complex, not least because of sample heterogeneity, but the dominant one in the PSIIm-S/27 sample appears to involve a modification of the Q_B_ site, affecting its proton-coupled electron transfer properties and its exchange with the PQ pool. Another important feature of this complex is the diminished affinity for bicarbonate and the significant positive redox shift of the Q_A_/Q_A_^•−^ reduction potential that appears to be present when the bicarbonate is not bound. Just such a shift occurs in standard granal PSII dimers when bicarbonate is released upon Q_A_^-^• accumulation (Brinkert et al., 2016). The redox shift protects against the well-characterised photodamage arising from chlorophyll triplet-mediated, singlet-oxygen generation (Johnson et al., 1995). This kind of protection is expected to be important in a near-intact PSII that is switched-off in transit, either after photoactivation or prior to repair.

## Material and Methods

### Growth and cultivation of tobacco plants

Transplastomic plants of *Nicotiana tabacum*, which have a hexa-histidine tag sequence at the 5 end of the gene coding for the PsbE subunit were used for this work (Fey et al., 2008). Plants were kept at a constant temperature of 25°C, at 50% relative humidity, and grown for 10–12 weeks under a light regime of 12 h/day, with a light intensity of 150–200 μmol photons s^-1^ m^-2^.

### Thylakoids preparation and PSII core solubilizations

Thylakoid membranes and PSII cores were prepared as previously reported (Haniewicz *et al*., 2013; Haniewicz *et al*., 2015), with only minimal modifications in the solubilization step. Briefly, PSII core complexes retaining the subunits PsbS and Psb27 were obtained from thylakoid membranes solubilized for 5 min at 4°C at a final chlorophyll concentration of 3 mg/mL. After solubilization, the unsolubilized fraction was separated by centrifugation at 35000 g for 10 min at 4°C. The unsolubilized fraction underwent a second solubilization step to isolate the PSII core complexes lacking PsbS and Psb27. This second solubilization took place for 15 min at 4°C at a final chlorophyll concentration of 1 mg/mL. Also, after this second solubilization, the unsolubilized fraction was separated by centrifugation at 35000 g for 10 min at 4°C. In both cases, the solubilization was carried out in the dark, adjusting the chlorophyll concentration with Grinding buffer (20 mM MES–NaOH, pH 6.5, 5 mM MgCl_2_) and using 20 mM (1.02 %) n-Dodecyl-β-D-maltoside (β-DDM).

### PSII core complexes isolation

Photosystem II samples were prepared using Ni-affinity chromatography and a subsequent step of size exclusion chromatography as reported in Haniewicz *et al*. (2013) for PSII complexes retaining the subunit PsbS, and according to Haniewicz *et al*. (2015) for PSII complex lacking the subunit PsbS. For the size exclusion chromatography step, buffer containing 20 mM MES–NaOH, pH 6.5, 5 mM MgCl_2_ and with 0.02% (~0.39 mM) β-DDM (buffer A). Previously a slightly higher detergent content was used of 0.03% (~ 0.59 mM) (Haniewicz *et al*., 2013; Haniewicz *et al*., 2015). In these studies, all chromatography columns were subjected to the ReGenFix procedure (https://www.regenfix.eu/) for regeneration and calibration prior use.

### Polyacrylamide gel electrophoresis

Denaturing Sodium Dodecyl Sulphate-Polyacrylamide Gel Electrophoresis (SDS-PAGE) consisted in 10% (w/v) separating polyacrylamide/urea gels with 4% (w/v) stacking gels (Piano et al., 2010; Collu et al., 2017). Samples were denatured with Rotiload (Roth) at room temperature before loading, and, after electrophoresis gels were stained with Coomassie brilliant blue G250.

### Absorption, CD spectroscopy and chlorophyll determination

The protein content of thylakoids was assessed through three independent measurements based on the concentrations of Chl *a* and Chl *b*. The absorption of chlorophylls extracted in 80% (v/v) acetone, in a dilution factor of 200 or 500, was measured with a Pharmacia Biotech Ultrospec 4000 spectrophotometer, and their relative concentrations were calculated according to Porra *et al*. (1989). CD spectra were the average of three accumulations recorded at a sensitivity of 100 mdeg and a scan speed of 100 nm/min using a CD spectrometer JASCO J-810. Absorption and CD spectra were recorded at room temperature in the range of 370-750 nm, with an optical path length of 1 cm and a band-pass of 2 nm. Spectra were recorded on an absorption Ultra Micro quartz cell with 10 mm light path (Hellma Analytics). In all cases, measurements were performed in a range between 0.01 and 0.2 mg/mL Chls, and samples were diluted in buffer A.

### Fluorescence spectroscopy and kinetics

Emission and excitation spectra were recorded on a Jasco FP-8200 spectrofluorometer at 4°C in 0.1 nm steps and 3 nm band-pass. Spectra were corrected for the photomultiplier sensitivity using a calibrated lamp spectrum. Emission spectra in the range of 600-750 nm were recorded using the main absorption bands as excitation wavelength (437 nm in Fig 2). Fluorescence spectra were recorded on a fluorescence Ultra Micro quartz cell with 3 mm light path (Hellma Analytics). The flash-induced increase and the subsequent decay of chlorophyll fluorescence yield and the values of F_0_, F_m_ and F_v_ were measured with a fast double modulation fluorimeter (FL 3000, PSI, Czech Republic). The sample concentration was 5 μg Chl/ml in buffer A. Samples were subjected to a pre-illumination in room light for 10 seconds followed by a period of 5 to 10 minutes of dark adaptation.

Multicomponent deconvolution of the measured curves was done by using a fitting function with three components based on the widely used model of the two-electron gate (Croft and Wraight, 1983; Vass et al., 1999). The fast and middle phases were simulated with exponential components. However, slow recombination of Q_A_^•−^ via charge recombination has been shown to obey hyperbolic kinetics corresponding to an apparently second order process (Bennoun, 1994), most probably the result of stretched exponentials indicative of inhomogeneity in this time-range. Therefore, the data were fitted with a linear combination of two exponentials and a hyperbolic component, where F_(t)_ is the variable fluorescence yield, F_0_ is the basic fluorescence level before the flash, A_1_–A_3_ are the amplitudes, T_1_–T_3_ are the time-constants, from which the half-lives were calculated via t_(1/2)_=ln 2·T for the exponential components, and t_(1/2)_ is the T for the hyperbolic component.

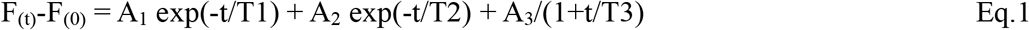

When the herbicide 3-(3,4-dichlorophenyl)-1,1-dimethylurea (DCMU) was added, 10 μM of an ethanolic solution were added in the dark to 1 mL of protein solution in buffer A, prior to the 5 minutes dark adaptation step.

The kinetics measured as a function of increasing concentrations of bicarbonate (0.5, 1, 5, 10 mM) were fitted with a hyperbolic curve (Eq. 2) from which the apparent dissociation constant (K_d app_) was calculated.

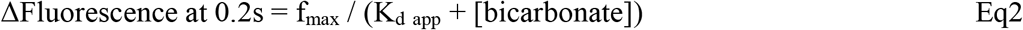

where ΔFluorescence at 0.2 s is the difference in fluorescence value at 0.2 s subtracted of the value before any addition of bicarbonate, and f_max_ is the fluorescence difference value when the binding site is fully occupied. The pH was monitored upon addition of bicarbonate to make sure that no shifts in pH occurred.

### Oxygen evolution

The oxygen evolution was measured with a Clark-type electrode (Hansatech, England) at 20°C, with 1 mM 2,6-dichloro-p-benzoquinone and 1 mM ferricyanide as electron acceptors in the reaction mixture. Measurements were carried on samples with a chlorophyll concentration of 1 mg/mL diluted 20 times in buffer A to a final concentration of 50 μg Chl/mL. Three independent measurements were done on the same preparation to test the activity. Reactions were started with illumination from a white light source (400-700 nm) with a Photosynthetically Active Photon Flux Density (PPFD) of 700-800 μmoles of photons m^-2^ s^-1^. For the effect of bicarbonate on PSIIm-S/27 samples, 5 mM NaHCO_3_ was added to the reaction mixture.

### Thermoluminescence

Thermoluminescence (TL) was measured with a lab-built apparatus, essentially as described in (Ducruet and Vass, 2009) but using a GaAsP photomultiplier H10769A-50 (Hamamatsu). Samples were pre-illuminated with room light (~ 20 μmol m^-2^ s^-1^) for 10 s, dark-adapted for 5 to 10 min and then cooled to 5°C. After 2 min, samples were excited with a single turnover saturating flash. Finally, samples were rapidly cooled to −15°C and luminescence was recorded with a 20°C min^-1^ heating rate. The sample concentration was 5 μg Chl/ml in buffer A.

## Acknowledgments

This work was carried out with the support of the program Homing Plus (Foundation for Polish Science) grant No: 2012-6/10 co-financed by the European Union under the European Regional Development Funds (to D.P.), the PRELUDIUM programme (National Science Centre) grant number DEC-2012/05/05/N/NZ1/01922 (to P.H.), Biotechnology and Biological Sciences Research Council (BBSRC) Grants BB/K002627/1 and BB/R00921X (to A.W.R.). We thank J-M Ducruet for help with setting up the thermoluminescence.

## Author contributions

A.F., A.W.R., D.P., and C.B. designed research; A.F., K.P., D.P., P.H., and D.F. performed research; D.P. and D.F. contributed new reagents/analytic tools; A.F., K.P., A.W.R., C.B., M.B., and D.P. analyzed data; A.F. A.W.R., D.P., P.H., D.F., C.B., M.B., and M.C.L. wrote the paper.

